# Microdomain protein Nce102 is a local sensor of plasma membrane sphingolipid balance

**DOI:** 10.1101/2021.12.06.471370

**Authors:** Jakub Zahumenský, Caroline Mota Fernandes, Petra Veselá, Maurizio Del Poeta, James B. Konopka, Jan Malínský

## Abstract

Sphingolipids are essential building blocks of eukaryotic membranes and important signalling molecules, tightly regulated in response to environmental and physiological inputs. Mechanism of sphingolipid level perception at the plasma membrane remains unclear. In *Saccharomyces cerevisiae*, Nce102 protein has been proposed to function as sphingolipid sensor as it changes its plasma membrane distribution in response to sphingolipid biosynthesis inhibition. We show that Nce102 redistributes specifically in regions of increased sphingolipid demand, e.g., membranes of nascent buds. Furthermore, we report that production of Nce102 increases following sphingolipid biosynthesis inhibition and Nce102 is internalized when excess sphingolipid precursors are supplied. This suggests that the total amount of Nce102 in the plasma membrane is a measure of the current need for sphingolipids, whereas its local distribution marks sites of high sphingolipid demand. Physiological role of Nce102 in regulation of sphingolipid synthesis is demonstrated by mass spectrometry analysis showing reduced levels of complex sphingolipids and long-chain bases in *nce102*Δ deletion mutant. Nce102 behaves analogously in human fungal pathogen *Candida albicans*, suggesting a conserved principle of local sphingolipid control across species.

## INTRODUCTION

Cellular membranes are laterally segmented into functionally distinct microdomains with specific composition and morphology. One of the best-characterised microdomains in the plasma membrane of eukaryotes is the <u>m</u>embrane <u>c</u>ompartment of <u>C</u>an1 (MCC; Malínská *et al*, 2003). The MCC accumulates a range of integral proteins, including nutrient importers and tetraspan proteins of Sur7 and Nce102 families (Young *et al*, 2002; Malinska *et al*, 2004; Grossmann *et al*, 2007, 2008; Fröhlich *et al*, 2009; Bianchi *et al*, 2018; Busto *et al*, 2018). It is believed to be rich in ergosterol based on local accumulation of the filipin dye (Grossmann *et al*, 2007). The MCC has a typical furrow shape (Stradalova *et al*, 2009) that is stabilized on the cytosolic side of the plasma membrane by the eisosome, a large hemitubular protein scaffold made of BAR (*Bin/Amphiphisin/Rvs*) domain-containing proteins Pil1 and Lsp1 that directly bind to PI(4,5)P_2_ (phosphatidylinositol 4,5-bisphosphate) in the plasma membrane and induce membrane curvature (Zhang *et al*, 2004; Walther *et al*, 2006; Stradalova *et al*, 2009; Ziółkowska *et al*, 2011; Karotki *et al*, 2011; Olivera-Couto *et al*, 2011; Kabeche *et al*, 2011, 2014).

The integral plasma membrane protein Nce102 is a structurally and functionally important component of the MCC/eisosome. Its presence in the MCC is essential not only for the accumulation of nutrient transporters in the microdomain (Grossmann *et al*, 2008) but also for the stability of the MCC/eisosome structure as a whole. Deletion of *NCE102* leads to a significant decrease in the number of MCC/eisosome microdomains in the plasma membrane of *S. cerevisiae* (∼50%; Fröhlich *et al*, 2009) and *S. pombe* (*fhn1*Δ, the sole *NCE102* homologue in *S. pombe*; ∼15%; Kabeche *et al*, 2011), and in the germling heads of *A. nidulans* (∼30%; Athanasopoulos *et al*, 2015). The MCC domains that are formed in the *nce102*Δ mutant lack their typical furrow shape and are flat (Stradalova *et al*, 2009). Interestingly, deletion of as few as 6 amino acid residues from the C-terminus of Nce102 is sufficient to abolish Nce102 accumulation in the MCC, resulting in a phenotype comparable with deletion of the whole gene in respect to sequestration of nutrient transporters in the microdomain as well as its morphology (Loibl *et al*, 2010).

While the physiological function of Nce102 is still largely unveiled, it has been previously shown in *S. cerevisiae* that the protein leaves the MCC following inhibition of sphingolipid biosynthesis in a growing culture, induced either by myriocin or aureobasidin A. Based on this result the authors suggested that Nce102 might function as a sphingolipid sensor (Fröhlich *et al*, 2009). Sphingolipids, especially inositolphosphoceramides and their mannosylated derivatives are very abundant in the fungal membranes (Kübler *et al*, 1996; Bagnat *et al*, 2000), representing the majority of sugar-containing lipids in *S. cerevisiae* (Souza & Pichler, 2007). Besides their essential structural function, sphingolipids and their precursors (ceramides and long-<u>c</u>hain <u>b</u>ases – dihydrosphingosine and phytosphingosine; LCBs) are important signalling molecules in a wide range of cellular functions, including response to environmental and endoplasmic reticulum stress (Dickson *et al*, 1997; Cowart *et al*, 2003; Jenkins, 2003; Dickson, 2010; Piña *et al*, 2018). It is therefore crucial that their levels be tightly regulated. The current working model is that the localization of Nce102 within the plasma membrane responds to the level of complex sphingolipids. Dissipation of Nce102 out of the MCC in response to inhibition of sphingolipid biosynthesis results in hyperphosphorylation of the core eisosome component Pil1, and in turn disassembly of the eisosome hemitubule structure (Fröhlich *et al*, 2009). This leads to the release of Slm1,2 proteins from the MCC/eisosome that activate Target of Rapamycin Complex 2 (TORC2; Daquinag *et al*, 2007; Berchtold *et al*, 2012; Riggi *et al*, 2020), the master lipid biosynthesis regulator (reviewed in Zahumensky & Malinsky, 2019). While considerable attention has been devoted to the steps leading to activation of TORC2 and in turn sphingolipid biosynthesis, the molecular details of how and why Nce102 leaves the MCC upon sphingolipid depletion remain unresolved.

In this paper, we report that the redistribution of Nce102 out of the MCC is a marker of locally increased sphingolipid demand. We show that following inhibition of sphingolipid biosynthesis, active progression through the cell cycle is necessary for plasma membrane sphingolipid levels to decrease and propose that budding of new cells is the mechanism responsible. In addition, we show an increase in the level of Nce102 in response to elevated sphingolipid demand, possibly to increase TORC2 activity, and internalization of Nce102 to the vacuolar membrane when excess sphingolipid precursors are supplied. Consistently, the *nce102*Δ deletion strain has decreased levels of complex sphingolipids relative to the isogenic wild type. We also report that Nce102 of the prominent human fungal pathogen *Candida albicans* behaves in a manner analogous to Nce102 in *S. cerevisiae*.

## RESULTS AND DISCUSSION

### The amount of Nce102 in the plasma membrane reflects the current sphingolipid demand

The inhibition of sphingolipid biosynthesis by the addition of myriocin to a growing *S. cerevisiae* culture leads to dissipation of Nce102 out of the MCC into the surrounding plasma membrane (Fröhlich *et al*, 2009; Fig 1A, C). Investigating this phenomenon further, we noticed that the change in Nce102 localization was accompanied by an ∼1.5-fold increase in mean cellular Nce102-GFP fluorescence intensity (Fig 1A, D). Mean intensity of the plasma membrane and the cell interior increased proportionally (Fig EV1). In addition, GFP-stained intracellular structures emerged that we identified as the endoplasmic reticulum (Fig EV1). These results suggested that inhibition of sphingolipid synthesis triggered *de novo* synthesis of Nce102, which we confirmed by western blotting after myriocin treatment (Fig 1E). Consistent with this, but contrary to results reported previously (Fröhlich *et al*, 2009), we found that Nce102-GFP dissipation out of the MCC in response to myriocin treatment was dependent on active *de novo* protein synthesis. When we simultaneously blocked *de novo* protein synthesis with cycloheximide and sphingolipid biosynthesis with myriocin, the myriocin effect on Nce102 localization and amount was completely abolished (Fig 1A, C, D, E). Analogous behaviour was observed previously for the eisosome core component Pil1 (Zhang *et al*, 2004).

**Figure 1.**
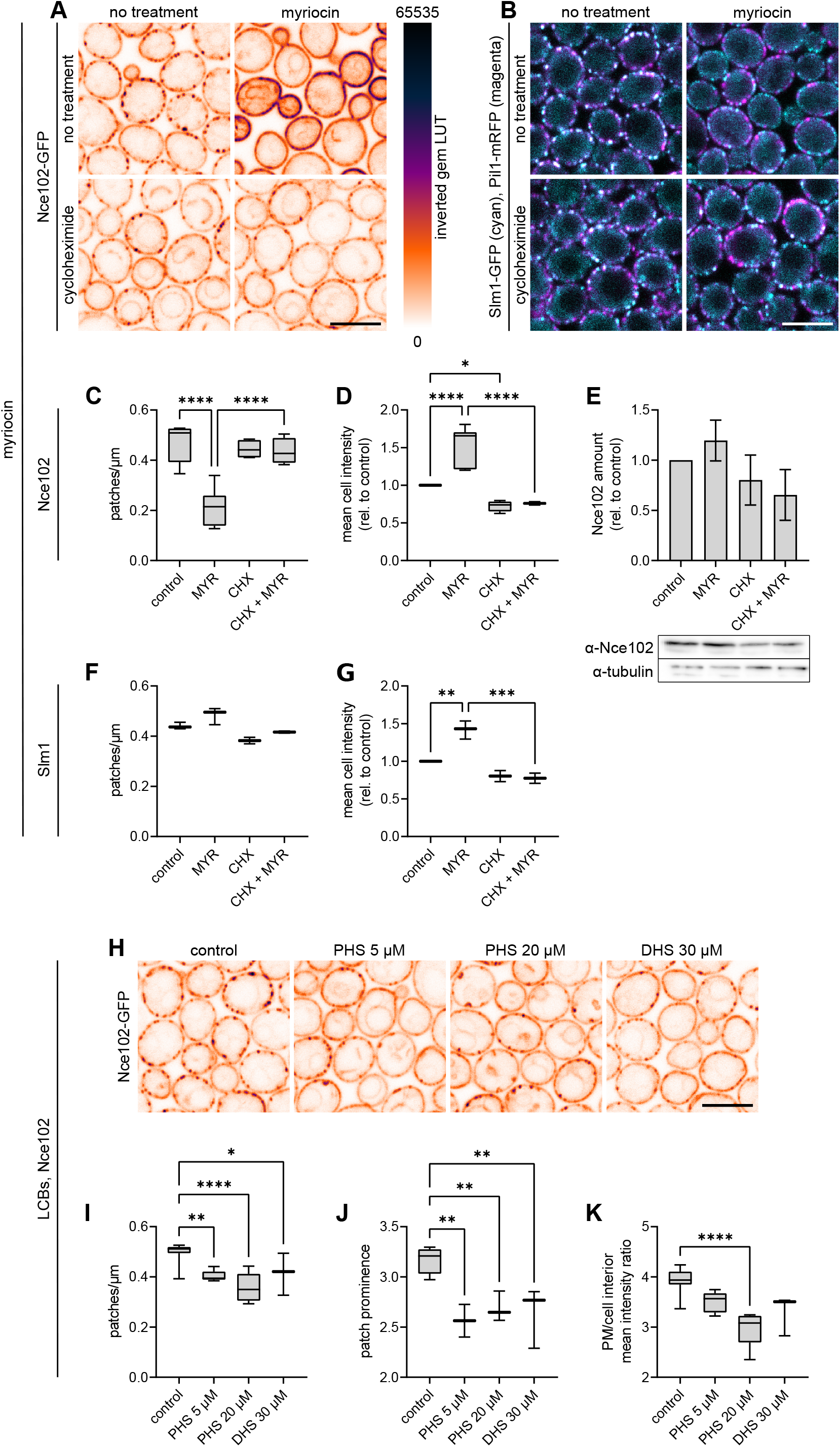
Inhibition of sphingolipid biosynthesis induces expression of Nce102 and Slm1. A, B, H Confocal microscopy images of *S. cerevisiae* cells expressing either *NCE102-GFP* (A, H) or *SLM1-GFP* (cyan) together with *PIL1-mRFP* (magenta)(B) cultivated for 6 hours and exposed to indicated chemicals for 2 hours. Scale bars: 5 μm. C, D Quantification of the number of Nce102-GFP patches per μm of cell cross-section circumference (C) and mean cell GFP intensity (D) in cultures treated as in (A). 4-7 biological replicates (n = 170–230 cells in each condition). E Western blot quantification of Nce102 amount in cultures treated as in (A). Data are presented as mean ± SD from 3 biological replicates. The results did not reach statistical significance (one-way ANOVA) because of variability between experiments. However, the Nce102 amount was always higher in myriocin-treated cultures than in the control in three independent experiments. Representative membrane with tubulin as loading control. F, G Quantification of the number of Slm1-GFP patches per μm of cell cross-section circumference (F) and mean cell GFP intensity (G) in cultures treated as in (B). 3 biological replicates (n = 150– 200 cells in each condition). I–K Quantification of the number of Nce102-GFP patches per μm of cell cross-section circumference (I), patch prominence, i.e., relative GFP intensity in patches vs surrounding plasma membrane (J) and the ratio of mean GFP fluorescence intensity in the plasma membrane (PM) and the cell interior (K) in cell cultures treated as in (H). 3 biological replicates (n = 300–400 cells in each condition). Data information: myriocin (MYR) – 10 μM, cycloheximide (CHX) – 100 μg/ml, LCBs – long-chain bases, DHS – dihydrosphingosine, PHS – phytosphingosine. In (A, H), LUT – inverted gem (depicted on the right-hand side of Fig 1A). In (C, D, F, G, I, J, K) data are presented as box plots. Bars represent the median. Error bars range from min to max value. * – P ≤ 0.05; ** – P ≤ 0.01; *** – P ≤ 0.001; **** – P ≤ 0.0001. One-way ANOVA. No statistically significant difference between the treatments was found in (F).

According to the current model, the dissipation of Nce102 out of the MCC destabilizes the whole MCC/eisosome leading to release of Slm1,2 proteins from the microdomain and in turn activation of TORC2 and sphingolipid biosynthesis (Tabuchi *et al*, 2006; Berchtold *et al*, 2012; Niles *et al*, 2012). To verify this, we monitored changes in Slm1-GFP localization following treatment of cells with myriocin. Indeed, there was strong Slm1-GFP signal dissipation and a decrease in overlap of Slm1-containing membrane patches with Pil1-mRFP signal (Fig 1B), consistent with previously published results (Berchtold *et al*, 2012; Busto *et al*, 2018). Analogous to Nce102 the mean cellular Slm1-GFP intensity increased following sphingolipid biosynthesis inhibition. When we inhibited *de novo* protein synthesis by cycloheximide Slm1-GFP retained its colocalization with Pil1-mRFP and change of mean cellular Slm1-GFP intensity was again abolished (Fig 1F, G).

We were also wondering about the behaviour of the proposed sphingolipid sensor under conditions of elevated sphingolipid amount. To achieve this state, we supplied the cells with surplus dihydrosphingosine (DHS) or phytosphingosine (PHS), long-chain bases used by the cells as sphingolipid precursors. 2-hour treatment of exponential cells with either PHS or DHS resulted in partial dissipation of Nce102-GFP patches and emergence of intracellular structures strongly stained with GFP (Fig 1H–K). We identified these as vacuoles by comparison with bright field images (Fig EV1). These data indicate that in the presence of surplus sphingolipid precursors, Nce102-GFP redistributes from the MCC into the surrounding plasma membrane and in turn to the vacuolar membrane.

### Nce102 dissipation out of the MCC in response to sphingolipid inhibition is dependent on active budding

We examined in more detail the necessity of active *de novo* protein synthesis for dissipation of Nce102 out of the MCC in response to sphingolipid biosynthesis inhibition. Among other effects, cycloheximide-induced block of protein synthesis ultimately leads to cell cycle arrest (Popolo *et al*, 1982). As progression through the cell cycle is tightly linked to the sphingolipid levels (Montefusco *et al*, 2014; Chauhan *et al*, 2016), we hypothesized that the dissipation of Nce102 out of the MCC upon myriocin treatment required active cell cycle progression.

First, we considered that myriocin is not growth-inhibitory within the time frame of our experiments (Sun *et al*, 2000). Further, in our timelapse experiments, it took ∼80 minutes for the Nce102-GFP patch density along the plasma membrane to drop to one half (Fig 2B, Movie EV2), which was sufficient for cells with small buds at t=0 min to finish cytokinesis and for newly formed buds to grow almost to full size. As the buds increased in size, the patchy pattern gradually vanished in their respective mother cells, while Nce102-labelled MCC patches were not formed at all in the growing daughters (Fig 2A). Cells that did not bud within the time frame of the experiment (120 min) retained the patchy Nce102-GFP distribution along the plasma membrane (note that due to the use of confocal microscopy for the detection of fluorescently labelled Nce102-GFP, not all buds seen in the bright field image are visible in the fluorescence channel showing just a single transversal cell section; compare Movies EV2 and EV3).

**Figure 2.**
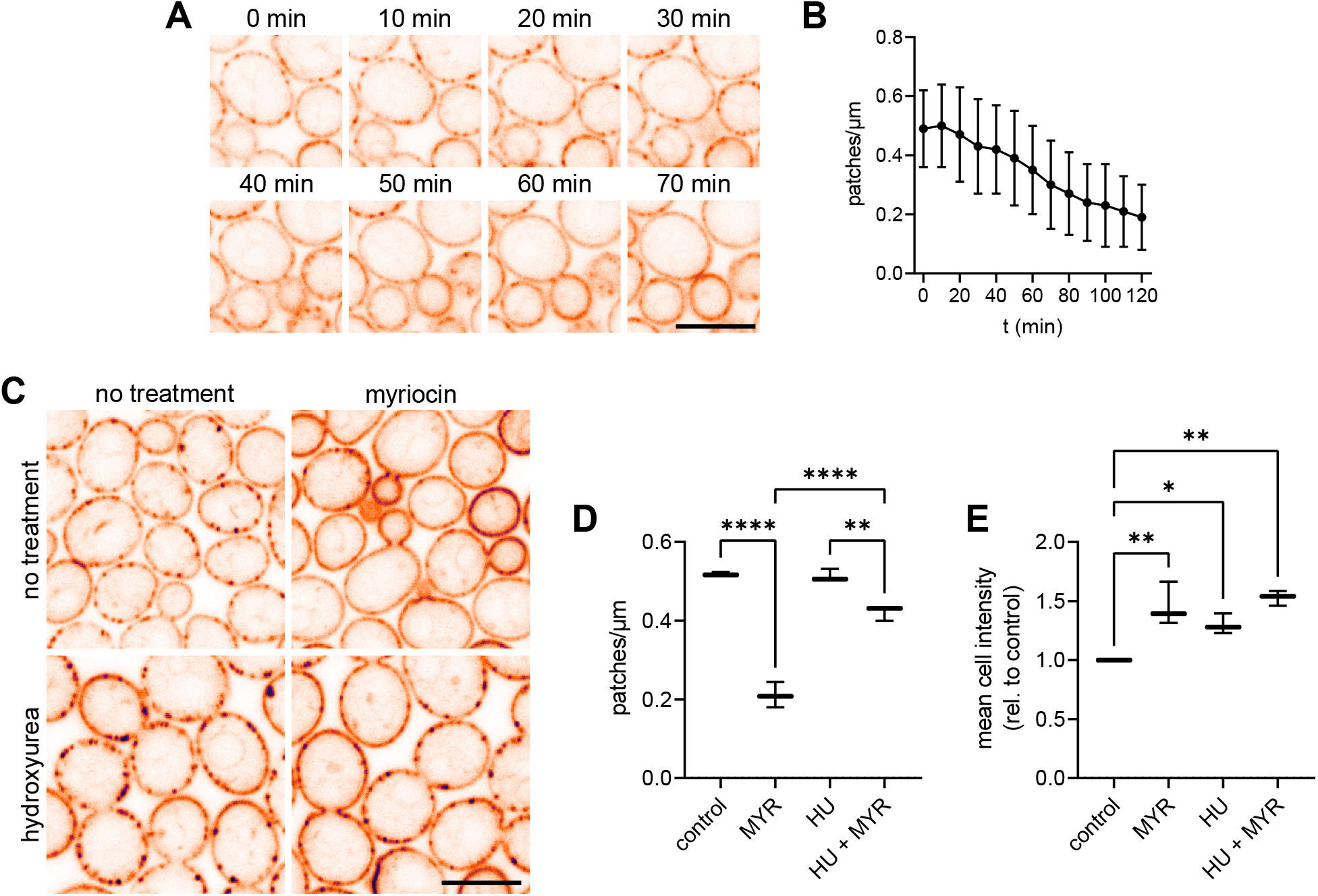
Nce102 dissipation out of the MCC in response to sphingolipid inhibition is dependent on active budding. A Confocal microscopy images of *S. cerevisiae* cells expressing *NCE102-GFP* cultivated for 6 hours, treated with myriocin and imaged in a time-lapse manner. B The dependence of Nce102-GFP patch density along the plasma membrane (number of patches per μm of cell cross-section circumference) on time quantified in images from a single time-lapse experiment (Movie EV2). Data are presented as mean ± SD from n = 150 cells in each image. C Confocal microscopy images of *S. cerevisiae* cells expressing *NCE102-GFP* cultivated for 5 hours, treated with hydroxyurea for 3 hours and subsequently for 2 hours with myriocin. D, E Quantification of the number of Nce102-GFP patches per μm of cell cross-section circumference (D) and mean cell intensity (E) in cultures treated as in (C). Data are presented as box plots. Bars represent the median. Error bars range from min to max value. 3 biological replicates (n = 150–200 cells in each condition). Data information: In (A,B), myriocin was added at t=0 min. In (C-E) myriocin (MYR) – 10 μM, hydroxyurea (HU) – 200 mM. In (A,C), Scale bars: 5 μm. LUT – inverted gem (depicted on the right-hand side of Fig 1A). In (D,E), * – P ≤ 0.05; ** – P ≤ 0.01; **** – P ≤ 0.0001. One-way ANOVA.

To directly test whether an active cell cycle is indeed required for the myriocin-induced change in Nce102 localization and abundance, we treated exponentially growing cells with 200 mM hydroxyurea for 3 hours before inhibiting sphingolipid biosynthesis by 2-hour exposure to myriocin. At the used concentration, hydroxyurea completely inhibits DNA synthesis resulting in cell cycle arrest in the S-phase. At the same time, *de novo* protein synthesis is decreased by only ∼30% (Slater, 1973; Rossio & Yoshida, 2011). As apparent from Fig 2C–E, arresting the cell cycle before myriocin addition strongly prevented changes in both localization and abundance of Nce102. Based on our results we conclude that cell cycle progression is required for Nce102 dissipation out of the MCC in response to inhibition of sphingolipid biosynthesis.

While a decrease of cellular complex sphingolipids in response to myriocin treatment has been reported, the mechanism of this change has not been described. Even though sphingolipid biosynthesis is blocked, the cells continue to bud and grow normally for up to ∼5 hours (Sun *et al*, 2000), suggesting that pre-existing sphingolipids are used in the newly synthesized bud membranes. We propose that following myriocin treatment sphingolipids necessary for incorporation into the bud plasma membrane are rerouted to the growing bud from the mother cell, leading to a decrease of complex sphingolipid levels in its plasma membrane, triggering dissipation of Nce102 out of the MCC. As the septum at the bud neck constitutes a strong diffusion barrier (Takizawa *et al*, 2000) and myriocin addition inhibits endocytosis by lowering the amount of long-chain bases (Zanolari *et al*, 2000; Rispal *et al*, 2015), the transport of sphingolipids from the mother to the bud could be mediated by membrane contact sites between the plasma membrane and either endoplasmic reticulum and/or Golgi, and inheritance of endoplasmic reticulum and/or Golgi. This mechanism predicts the existence of a phospholipase C that would convert the complex sphingolipids in the outer leaflet of the plasma membrane into ceramide, which has been shown to readily flip-flop in large unilamellar vesicles (Pohl *et al*, 2009).

### The distribution of Nce102 reflects local demand for sphingolipids

In our microscopy images of exponentially growing cultures, Nce102-GFP distributed homogeneously within the plasma membrane of growing buds (Fig 1A, 2C, EV4) even in the absence of sphingolipid biosynthesis inhibitors. As the buds became larger, Nce102-GFP patches started to form, initially predominantly in areas close to the bud neck, where the membrane was already matured. This is consistent with a previous report showing that eisosomes (the cytoplasmic hemitubules directly interacting with the MCC) are formed only after the growing bud reaches a critical surface area of ∼20 µm^2^. As the bud grows further, more eisosomes are gradually formed, starting at the bud neck and progressing towards the bud tip (Moreira *et al*, 2009). Importantly, the MCC protein Sur7 in *C. albicans* behaves in an analogous fashion (Alvarez *et al*, 2008). The newly formed plasma membrane in the growing bud is rapidly expanding and presents an area with high sphingolipid demand, analogous to cellular plasma membrane under conditions of inhibition of sphingolipid biosynthesis. Nce102 is distributed homogeneously in both cases. We conclude that Nce102 is a local signal of increased sphingolipid demand when localized homogeneously in a particular area of the plasma membrane, i.e., outside of the MCC/eisosome. We have previously shown an analogous regulation of activity by sequestration into the MCC/eisosome for the exoribonuclease Xrn1 that is inactivated by binding to the eisosome in post-diauxic cultures (Vaškovičová *et al*, 2017).

According to the current model, dissipation of Nce102 out of MCC triggers eisosome disassembly and the release of TORC2 activators Slm1,2 (Daquinag *et al*, 2007; Berchtold *et al*, 2012; Riggi *et al*, 2020). It has been proposed that Nce102 itself could, together with another integral plasma membrane protein Sng1, physically interact with TORC2. These proteins were suggested to create a scaffold enhancing the interaction of Ypk1 with TORC2, increasing the effectiveness of sphingolipid biosynthesis activation via TORC2 → Ypk1 → SPT (serine palmitoyltransferase) pathway under such circumstances (García-Marqués *et al*, 2016). Physical interaction of Nce102 with Tor1 and Tor2 has indeed been reported previously (Aronova *et al*, 2007). The increase in Nce102 level in the plasma membrane that we report here should therefore lead to increased TORC2 activation and hence higher overall activity of the sphingolipid biosynthetic pathway. Consistently, overexpression of *NCE102* has been reported to prevent eisosome disassembly in response to myriocin treatment (Fröhlich *et al*, 2009). Supporting the presented line of reasoning further is the increase in abundance of the known TORC2 activator Slm1 following myriocin treatment, which we report here (Fig 1G). We also show that creating a state of surplus availability of sphingolipid precursors leads to internalization of Nce102 into the vacuolar membrane (Fig 1H), which could serve to prevent excessive activation of TORC2. We have shown recently ((Vaskovicova *et al*, 2020); also apparent below in Fig 4A) that in post diauxic cells, when the cellular sphingolipid levels are high, a part of the Nce102 population localizes to the ergosterol-rich domains of the vacuole. This might represent a pool of Nce102 “ready to be put into action” should the need arise for a fast increase in sphingolipid biosynthesis. Whether the Nce102 localized to the vacuolar membrane can be redirected back to the plasma membrane under such conditions remains to be verified.

### Mechanisms governing Nce102 sequestration in the MCC in response to sphingolipid levels

One question that remains to be answered is how Nce102 senses local sphingolipid levels. One option is that the Nce102 molecule changes conformation in response to a local drop in sphingolipid concentration. This is favoured by a previous study that analysed the flexible membrane topology of the Nce102 protein (Loibl *et al*, 2010). Analogous to the arginine importer Can1, conformational change in the Nce102 molecule could result in lower clustering of Nce102 in the curved membrane of the MCC microdomain (Gournas *et al*, 2018). In the case of Can1 and other MCC transporters, the conformational change is triggered by substrate binding (Gournas *et al*, 2018; Busto *et al*, 2018). For Nce102, another mechanism needs to be considered, for example, a post-translational protein modification. Since an intact C-terminal domain of Nce102 is essential for the protein’s accumulation in the MCC (Loibl *et al*, 2010), this area appears to be a good candidate in this respect. In fact, two recent high-throughput studies have identified T162 and S171 in the cytosolic C-terminal tail of Nce102 as potential phosphorylation sites (MacGilvray *et al*, 2020; Lanz *et al*, 2021). In the context of the results of this study, it is interesting to mention that it appears that the cyclin-dependent kinase Cdk1 might be involved in the phosphorylation of T162 in a cell cycle-dependent manner (Holt *et al*, 2009). The phosphorylation status of Nce102 could be regulated by long-chain bases, as is the case of the eisosome constituents Pil1 and Lsp1 (Zhang *et al*, 2004).

Here we report that not only lack but also a surplus of sphingolipid precursors, causes Nce102 to dissipate out of the MCC. Following the addition of PHS, the sphingolipid-to-ergosterol balance is severely affected, leading to free ergosterol deficiency. In such a situation, the MCC might serve as an ergosterol reservoir, providing the necessary interaction partners for the surplus sphingolipids. This change in the lipid landscape could result in redistribution of Nce102 out of the MCC. Once outside, Nce102 is endocytosed to the vacuolar membrane, where it appears to be stable. We previously reported internalization of Nce102 via multivesicular body endocytic pathway in post-diauxic cultures (Vaskovicova *et al*, 2020). It appears that Nce102 localizes to MCC under basal conditions to be protected from untimely endocytosis, analogous to nutrient transporters in the absence of their substrates (Grossmann *et al*, 2008; Gournas *et al*, 2018). Upon disturbance in the sphingolipid balance in either direction, Nce102 is released from the MCC and its level in the plasma membrane is tuned accordingly, either by *de novo* synthesis or endocytosis.

### Acute inhibition of ergosterol biosynthesis does not affect Nce102 localization

Preferential localization of Nce102 into ergosterol-enriched domains of cellular membranes (Grossmann *et al*, 2007; Vaskovicova *et al*, 2020) suggests that modifications of ergosterol levels could result in changes in the protein distribution. To test this, we blocked ergosterol biosynthesis by the addition of fluconazole, a potent inhibitor of the essential enzyme Erg11, which causes an 80% drop in ergosterol levels in 24-hour old *S. cerevisiae* cultures already at 1.5 μg/ml concentration (Demuyser *et al*, 2017). In *C. albicans* inhibition of Erg11 was shown to cause an increase in the synthesis of long-chain fatty acid-containing complex sphingolipids (Gao *et al*, 2018). Exposure of exponentially growing *S. cerevisiae* cells to 2 μg/ml fluconazole for 2 hours caused no significant change in either localization of Nce102-GFP or mean cellular GFP fluorescence (Fig 3).

**Figure 3.**
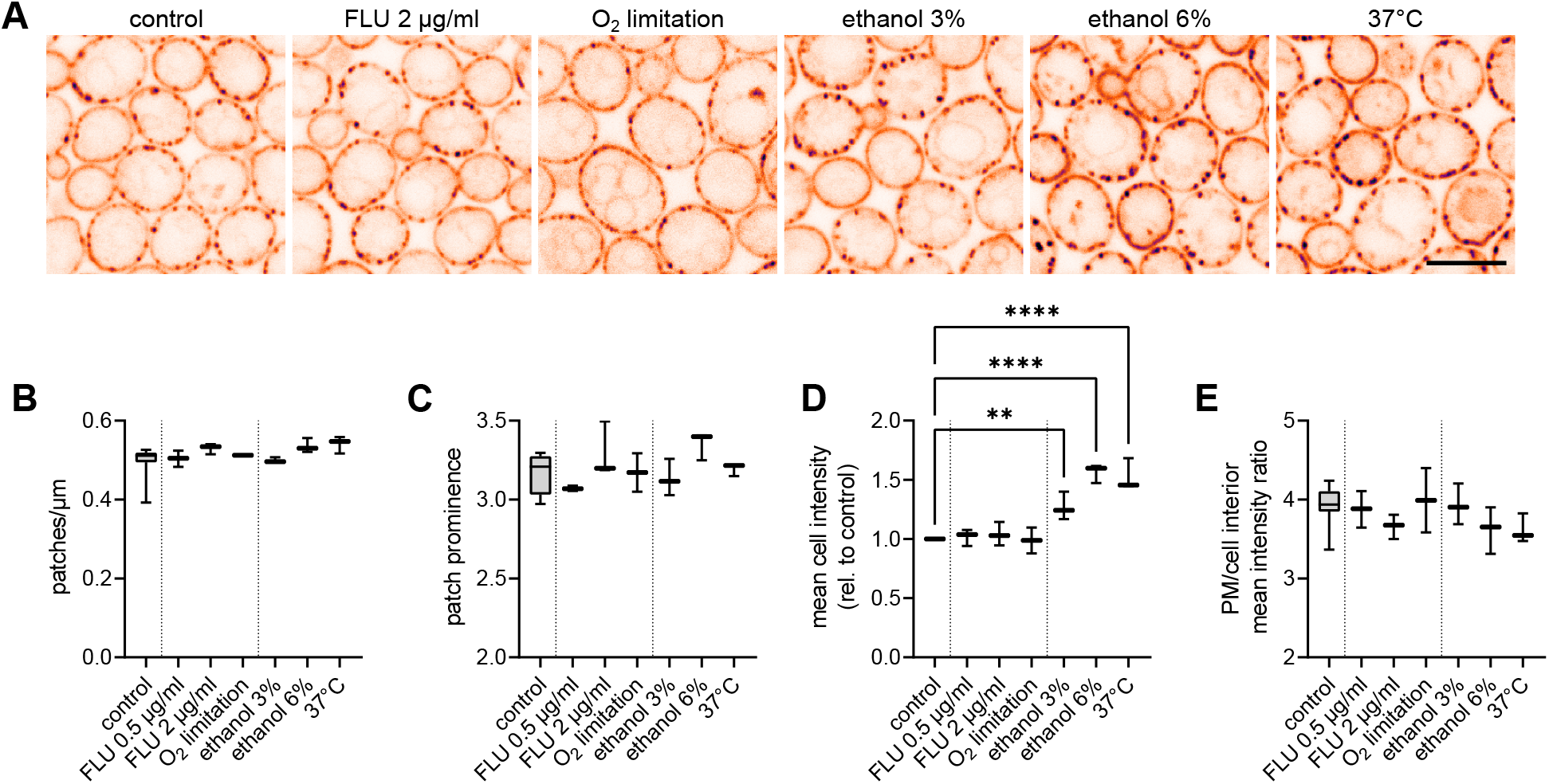
Nce102 localization and abundance change in response to increasing plasma membrane fluidity and sphingolipid levels. A Confocal microscopy images of *S. cerevisiae* cells expressing *NCE102-GFP* cultivated for 6 hours and exposed to indicated stress conditions for 2 hours. Scale bar: 5 μm. LUT – inverted gem (depicted on the right-hand side of Fig 1A). B–E Quantification of the number of Nce102-GFP patches per μm of cell cross-section circumference (B), patch prominence, i.e., relative GFP intensity in patches vs surrounding plasma membrane (C), mean cell intensity (D) and the ratio of mean GFP fluorescence intensity in the plasma membrane (PM) and the cell interior (E), in cultures grown for 6 hours and exposed to indicated stress conditions for 2 hours. Data are presented as box plots. Bars represent the median. Error bars range from min to max value. 3-5 biological replicates (n = 300–400 cells in each condition). In (D), ** – P ≤ 0.01; **** – P ≤ 0.0001. One-way ANOVA. No statistical difference was found between conditions in (B, C, E). Vertical dotted lines separate groups of stress conditions – left to right: control, ergosterol inhibition, increase of membrane fluidity. Data information: FLU – fluconazole

The ergosterol synthesis can also be inhibited by limiting the supply of oxygen, which is required for the production of heme, an ergosterol precursor (Jahnke & Klein, 1983). *S. cerevisiae* cultures grown exponentially in hypoxic conditions produce only ∼10% ergosterol of the amount made by their aerated counterparts (Gleason *et al*, 2011). In addition, lack of oxygen causes a decrease in plasma membrane fluidity due to defective desaturation of fatty acids (Sharma, 2006; Degreif *et al*, 2017). As after fluconazole treatment, we found no significant difference between Nce102 localization and abundance in cells exposed to oxygen-limiting conditions for 2 hours (Fig 3). We conclude that Nce102 distribution is not sensitive to acute changes in ergosterol levels.

### Acute increase in plasma membrane fluidity induces expression of Nce102

To increase the membrane fluidity of exponentially growing cells we either shifted the culture from optimal 28 to 37°C or added ethanol to the growth medium (Ingram, 1976; Hazel, 1995). The increase in membrane fluidity is normally compensated by an increase in biosynthesis of the rigidity-supplying complex sphingolipids (Jenkins *et al*, 1997; Wells *et al*, 1998; Aresta-Branco *et al*, 2011; Malinsky & Opekarová, 2016). Upon transferring of 28°C-grown exponential culture to 37°C for 2 hours, mean cellular Nce102-GFP intensity increased ∼1.5-fold (Fig 3A, D), indicating an increase in Nce102 protein level. Since the two treatments applied shift the plasma membrane fluidity in the same direction, we expected to find similar response of the proposed sphingolipid sensor Nce102 under these two conditions. Indeed, exposing exponential cells to ethanol caused an increase in the mean cellular fluorescence. The effect increased with ethanol concentration and at 6% was comparable with the shift to 37°C. We conclude that acutely increasing the fluidity of the plasma membrane triggers synthesis of Nce102.

### Nce102 localization and abundance changes following chronic stress

As we found that Nce102 plays a sensor role in the regulation of cellular response to acute sphingolipid imbalance and increase in plasma membrane fluidity, we decided to investigate if the protein is involved also in long-term adaptation to stress. Therefore, we applied the studies stress conditions for extended periods and monitored behaviour of Nce102. Cell cultures grown for 6 hours in the continuous presence of surplus ethanol and sphingolipid precursors, as well as under elevated temperature, exhibited characteristics comparable to exponential cells exposed to these stresses acutely for 2 hours (Fig 4A–D). Specifically, mean cellular GFP fluorescence increased, while GFP-stained intracellular structures became more prominent, especially following PHS addition. The excess of sphingolipid precursors also caused partial loss of Nce102 patches. Inhibition of ergosterol synthesis by either fluconazole addition or limiting oxygen supply had again no significant effect on Nce102.

**Figure 4.**
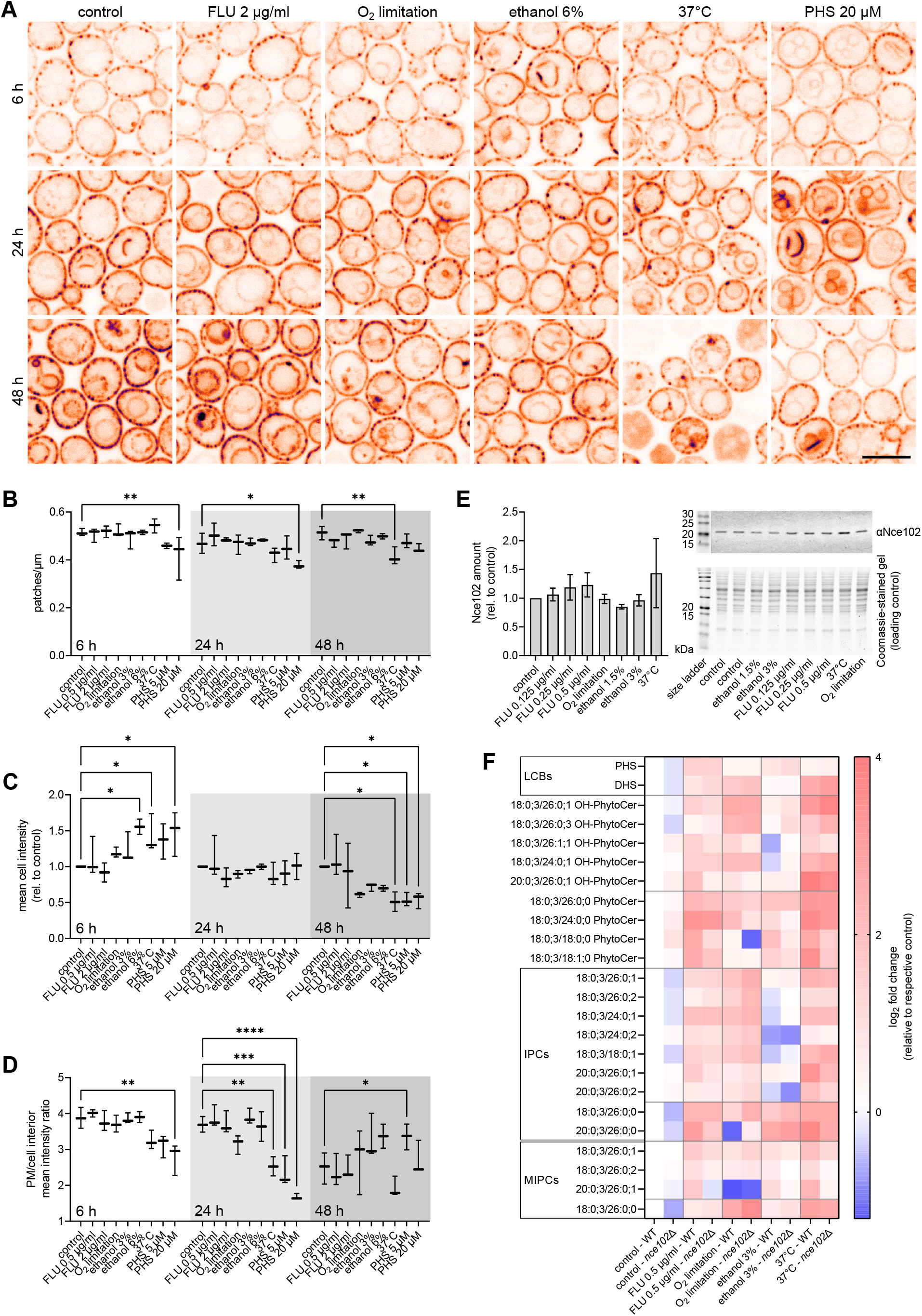
Nce102 response to chronic stress. A Confocal microscopy images of *S. cerevisiae* cells expressing *NCE102-GFP* treated with indicated stress upon inoculation and cultivated for the indicated time. Scale bars: 5 μm. LUT – inverted gem (depicted on the right-hand side of Fig 1A). B–D Quantification of the number of Nce102-GFP patches per μm of cell cross-section circumference (B), mean cell Nce102-GFP intensity (C) and the ratio of mean GFP fluorescence intensity in the plasma membrane (PM) and in the cell interior (D) in cell cultures grown for indicated time in the chronic presence of indicated stress conditions. 3 biological replicates (n = 150 cells in 48 h 37°C condition – dead cells were excluded from analysis; n = 250–400 cells in all other conditions). E Western blot quantification of Nce102 amount in *S. cerevisiae* cultures grown for 48 hours under chronic exposure to indicated stress. Data are presented as mean ± SD from 3 biological replicates. No statistically significant difference between the treatments was found (one-way ANOVA). Representative western blot membrane and Coomassie-stained gel used as loading control. Note the different order of treatments in the graph and on the gel/membrane. F Mass spectrometry analysis of differences in sphingolipid levels of *S. cerevisiae* wild type (WT) and *nce102*Δ treated with indicated stress conditions upon inoculation and cultivated for 48 hours. Values of stress-treated conditions relative to respective control; *nce102*Δ control relative to wild type control. 3 biological replicates. Data information: FLU – fluconazole, DHS – dihydrosphingosine, PHS – phytosphingosine, LCBs – long-chain bases, IPCs – inositol phosphoceramides, MIPCs – mannosyl inositol phosphoceramides. In (B, C, D), data are presented as box plots. Bars represent the median. Error bars range from min to max value. * – P ≤ 0.05; ** – P ≤ 0.01; *** – P ≤ 0.001; **** – P ≤ 0.0001. One-way ANOVA.

When we cultivated the cells for 24 hours either at elevated temperature or in the surplus of sphingolipid precursors the intracellular GFP fluorescence increased significantly. In the case of sphingolipid excess, this was accompanied by a decrease in the number of Nce102-GFP patches in the plasma membrane (Fig 4A, B, D). Exposing the cultures to either of these two stress conditions resulted in a decrease of mean cellular fluorescence and significant death of the cells (Fig 4A, C). The cultures cultivated in the presence of fluconazole, ethanol and under conditions of limited oxygen supply for 24–48 hours exhibited characteristics comparable to the control cells, suggesting possible adaptation of the growing cultures. According to biochemical analysis of *S. cerevisiae* cultures, 48-hour exposure to studied stress conditions did not result in a significant change in cellular Nce102 protein amount (Fig 4E; negative control in Fig EV5). We conclude that Nce102 plays a more significant role in the initial response to acute disruption of sphingolipid balance and increasing plasma membrane fluidity than in long-term adaptation to stress.

### Nce102 fine-tunes sphingolipid synthesis in response to chronic stress

We show above that the cellular localization and amount of Nce102 responds to changes in the sphingolipid composition and fluidity of the plasma membrane. As discussed above, this might be a tool for tuning TORC2 activity and in turn sphingolipid biosynthesis. To verify this, we analysed changes in cellular sphingolipid composition in response to the studied stress conditions and assessed the contribution of Nce102 protein to these changes.

As immediately apparent from Fig 4F (raw data in Appendix Table S1), deletion of *NCE102* caused a general decrease in the levels of long-chain bases (LCBs), inositolphosphoceramides (IPCs) and mannosyl inositolphosphoceramides (MIPCs), supporting the proposed role of Nce102 in the activation of sphingolipid biosynthesis. Interestingly, the levels of phytoceramides were generally higher in the deletion mutant, suggesting the possibility of a functional interplay between Nce102 and either IPC synthase (Aur1-Kei1) or Isc1, a yeast homologue of mammalian sphingomyelinases (Matmati & Hannun, 2008), responsible for conversion of complex sphingolipids back into ceramides.

As expected, all stress conditions resulted in an increase of sphingolipid levels across almost all detected classes, including long-chain bases. The wild type and the deletion mutant strain behaved similarly. The largest increase took place when cells were cultivated at 37°C, pointing to activation of *de novo* sphingolipid biosynthesis. While the degree of the stress-induced changes was not significantly different in the *nce102*Δ deletion mutant for most of the particular lipids, the hydroxylated phytoceramides as a group increased to a slightly greater extent in the *nce102*Δ mutant. The opposite was true for all other lipid classes.

Following cultivation with either ethanol or fluconazole, the non-hydroxylated phytoceramides and IPCs increased more extensively than other sphingolipids. In the case of ethanol, however, there was an overall decrease in hydroxylated IPC species suggesting a more general remodelling of the plasma membrane lipid landscape. The changes in either direction were again less extensive in the *nce102*Δ deletion mutant in response to both ethanol and fluconazole. Under conditions of limited oxygen supply, there is again an overall increase of the levels of all sphingolipid classes, except for a few particular lipids. While the increase in the amount of phytoceramides was less extensive in the absence of Nce102, all detected inositolphosphoceramides and mannosyl inositolphosphoceramides increased slightly more in the *nce102*Δ mutant in response to oxygen limitation.

Based on these data we conclude that Nce102 is involved not only in the regulation of the overall rate of sphingolipid biosynthesis (via regulation of TORC2 activation) but that it is functionally connected also to additional enzymes of the pathway. The resulting impact on the sphingolipid landscape depends on particular conditions and deserves to be studied in more detail.

### Nce102 behaviour is conserved in the human fungal pathogen *C. albicans*

As the MCC/eisosome microdomain is evolutionarily conserved throughout the fungal kingdom (Stradalova *et al*, 2009; Lee *et al*, 2015), we tested whether Nce102 exhibits the same behaviour also in the opportunistic human fungal pathogen *Candida albicans*. In healthy humans, *C. albicans* is a harmless inhabitant of the gastrointestinal tract (Neville *et al*, 2015). Therefore, cultures of this yeast grown at 37°C were used as controls. Just like in *S. cerevisiae*, 2-hour myriocin treatment of exponentially growing *C. albicans* cells leads to dissipation of Nce102-GFP from the MCC (Fig 5A, B), suggesting a conservation of the sensor function of Nce102. *C. albicans* cultures grown for 48 hours at 42°C exhibited large-scale dying of the cells, as did cultures chronically exposed to 6% ethanol and as low as 1 μg/ml fluconazole (Fig EV5). Therefore, we compared *Ca*Nce102-GFP localization and amount between cultures grown at optimal 37°C and suboptimal 30°C. Similarly, we used lower concentrations of ethanol and fluconazole. Both cultivation in the presence of 3% ethanol and at 30°C resulted in a higher Nce102-GFP patch density within the plasma membrane (Fig 5C, D), analogous to *S. cerevisiae* (compare with Fig 4A, B). This correlated well with the increased cellular amount of *Ca*Nce102-GFP (Fig 5F; negative control in Fig EV5). Inhibition of ergosterol synthesis by either fluconazole addition or limiting oxygen supply resulted in an increase of GFP intensity inside the vacuole (shifting the PM/cell interior GFP intensity ratio towards lower values), suggesting increased degradation of *Ca*Nce102-GFP (Fig 5C, E). Our data show that Nce102 behaves analogously in a laboratory *S. cerevisiae* strain and in a *C. albicans* strain derived from a clinical isolate, pointing to conservation of the sensing principle. The reported changes of Nce102 localization and level are more extensive in the pathogen, corresponding to its need to be able to react rapidly to changes in the environment caused by the immune system response of the host organism.

**Figure 5.**
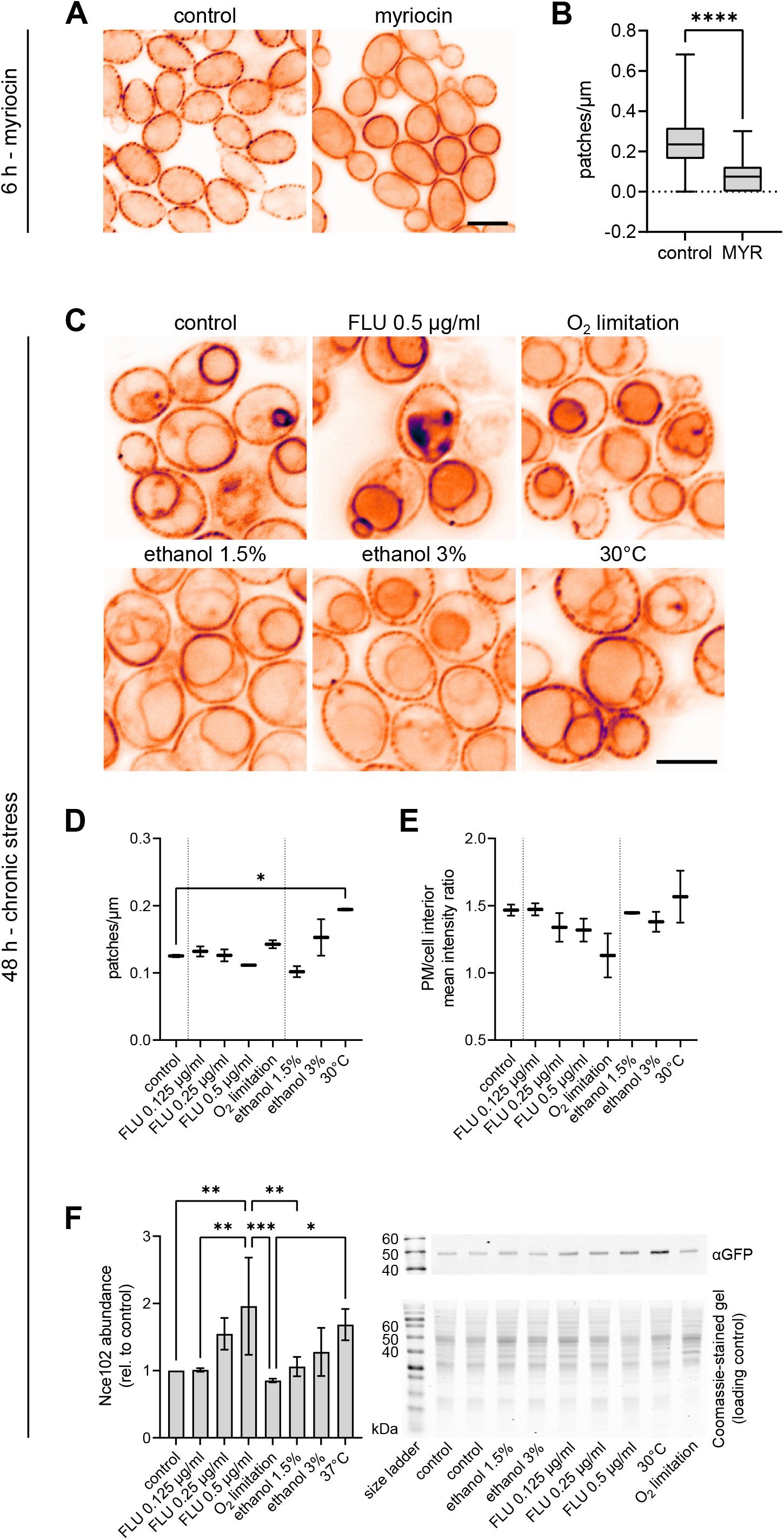
Nce102 behaves analogously in *C. albicans* and *S. cerevisiae*. A, C Deconvolved wide-field fluorescence microscopy images of *C. albicans* cells expressing *NCE102-GFP* cultivated for 6 hours and exposed to myriocin for 2 hours (A) or treated with indicated stress upon inoculation and cultivated for 48 hours (C). LUT – inverted gem (depicted on the right-hand side of Fig 1A). Scale bars: 5 μm. B, D Quantification of the number of *Ca*Nce102-GFP patches per μm of cell cross-section circumference following either 2-hour myriocin treatment (B) or chronic exposure to indicated stress (C). E Quantification of the ratio of mean GFP fluorescence intensity in the plasma membrane (PM) and the cell interior following chronic exposure to indicated stress. No statistically significant difference between the treatments was found (one-way ANOVA). F Western blot quantification of Nce102 amount in *C. albicans* cultures grown for 48 hours under chronic exposure to indicated stress. Data are presented as mean ± SD from 3 biological replicates. Representative western blot membrane and Coomassie-stained gel used as loading control. Note the different order of treatments in the graph and on the gel/membrane. Data information: myriocin (MYR) – 10 μM, FLU – fluconazole. In (B), one biological replicate (n = 975 cells in control and 512 cells in myriocin-treated culture). In (D,E), 2 biological replicates, n ≥ 100 cells in each condition). In (B,D,E,F), * – P ≤ 0.05; ** – P ≤ 0.01; *** – P ≤ 0.001; **** – P ≤ 0.0001. One-way ANOVA.

### Concluding remarks

In conclusion, our study provides important insights into the mechanisms by which the cell perceives the levels of sphingolipids in the plasma membrane and relays this information downstream. Specifically, we identified Nce102 as a local marker of increased sphingolipid demand within the plasma membrane. In addition, we show that the amount of the protein fine-tunes sphingolipid synthesis, probably via interaction with TORC2 and/or specific enzymes along the actual sphingolipid biosynthesis pathway. Uncovering the details of these interactions will be the objective of further studies. While disruptions of the MCC microdomain rarely lead to a growth-related phenotype in laboratory strains of *S. cerevisiae*, they are detrimental for morphogenesis and invasive growth of *C. albicans* (Douglas *et al*, 2012, 2013; Wang *et al*, 2016). Our study adds to the growing body of data indicating that the MCC/eisosome is a viable target in the treatment of fungal infections (Douglas & Konopka, 2019; Lanze *et al*, 2021).

## MATERIAL AND METHODS

### Strains and growth conditions

*S. cerevisiae* and *C. albicans* strains used in this study are listed in Table 1. The Y786 strain was constructed by genomic tagging of *PIL1* with *mRFP* in the Y549 strain using the YIp128-*PIL1-mRFP* integrative plasmid as described previously (Vaskovicova *et al*, 2015). Yeast cultures were cultivated in synthetic media (0.17% yeast nitrogen base without amino acids and ammonium sulphate, 0.5% ammonium sulphate and 2% dextrose) supplemented with required amino acids. In the case of *C. albicans*, 80 mg/l uridine was used instead of uracil. Unless stated otherwise, *S. cerevisiae* cultures were grown in an incubator equipped with an orbital shaker at 220 rpm at either 28°C (microscopy) or 30°C (western blot) and *C. albicans* in a tube roller at 37°C. Overnight cultures were diluted to OD_600_=0.2 in fresh media to start main cultures which were then incubated for desired time.

**Table 1:**
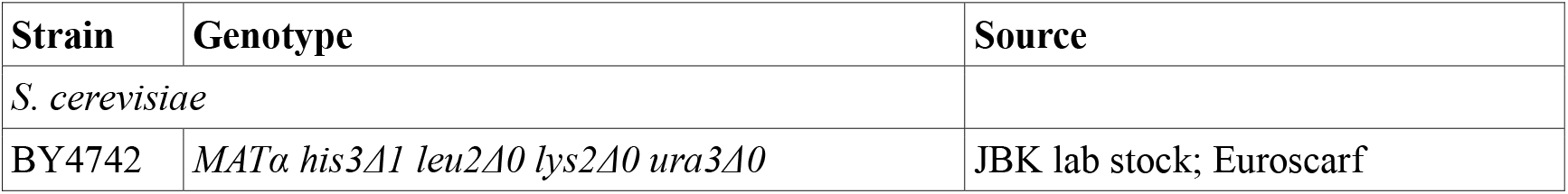
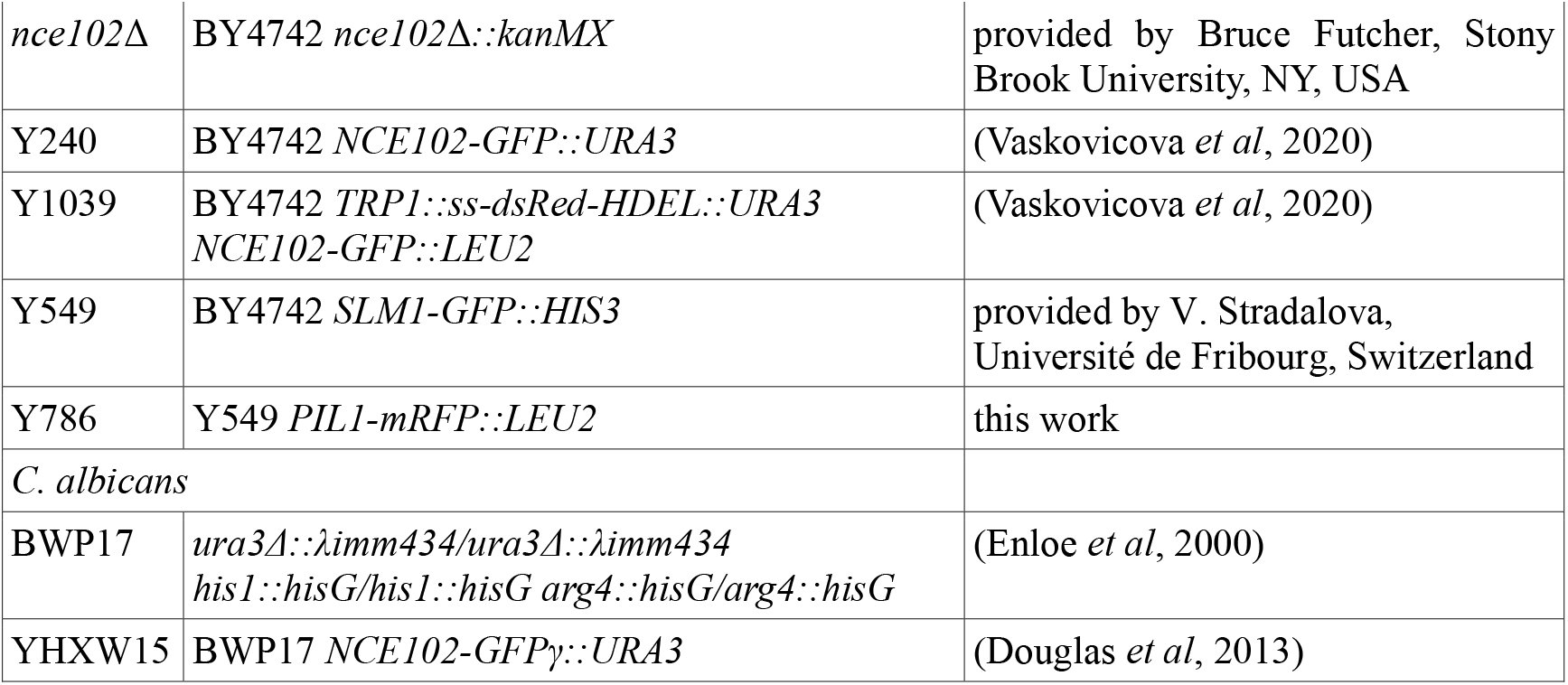
Strains used in the study

### Stress treatment

For assessment of the effects of acute stress, cultures were grown as described above for 6 hours, whereupon they were exposed to stress for 2 hours. For chemically induced stress, ethanol (96%), fluconazole (1 or 2 mg/ml ethanol stock), myriocin (2 mg/ml methanol stock), cycloheximide (10 mg/ml water stock), phytosphingosine (PHS, 5 mg/ml ethanol stock), dihydrosphingosine (DHS, 5 mg/ml methanol stock), hydroxyurea (2 M SC stock heated to 28°C prepared fresh before each experiment) were added to desired concentrations. All chemicals with exception of ethanol (Penta) were purchased from Merck. Oxygen limitation was induced by growing a larger volume of cell culture aerobically and then replacing 33 ml into a 50 ml Falcon tube that was closed tightly and sealed with parafilm to prevent the exchange of air with the surroundings. For heat stress, *S. cerevisiae* cells were shifted from 28 to 37°C; *C. albicans* was shifted from 37 to 30°C, as an increase to 42°C resulted in extensive dying of cells. Chronic stress was induced by the same agents applied when the growth of main cultures was started. These were then grown for 6, 24 or 48 hours and processed for microscopy, western blotting or lipid analysis as described in respective paragraphs below.

### Confocal fluorescence microscopy of live cells

Living yeast cells were grown as described above, either in the presence or absence of specific stress conditions and concentrated by brief centrifugation. A small amount of cell suspension was immobilized on a 0.17 mm cover glass by a thin film of 1% agarose prepared in 50 mM potassium phosphate buffer (pH 6.3). For time-lapse experiments of exponential cultures, the 1% agarose was prepared in a fresh SC medium to supplement nutrients. *S. cerevisiae* cells were imaged using a Zeiss LSM 880 and Olympus BX61WI scanning confocal microscopes. For *C. albicans* cells, 9 to 11-layered z-stacks were made using a Zeiss Axio Observer Z1/7 wide-field fluorescence microscope. All three microscopes were equipped with 100× PlanApochromat oil immersion objectives (NA=1.4). GFP and mCherry/dsRed were excited using a 488 nm Ar and a 559 nm LD laser, respectively. *C. albicans* images were drift-corrected and deconvolved in the proprietary Zeiss ZEN software, black edition. Image processing and analysis were performed in Fiji (ImageJ 1.53c) using custom-written macros (available at https://github.com/jakubzahumensky/Nce102_SL_paper) and cell segmentation masks made using Cellpose (Stringer *et al*, 2021). Statistical analysis was performed in GraphPad Prism 9 software.

### Western blot

For the analysis of Nce102 abundance in cells chronically exposed to stress, cultures (wild type BY4742 for *S. cerevisiae*; YHXW15 with GFP-tagged Nce102 for *C. albicans*) were grown for 48 hours in the absence or presence of indicated stress conditions and harvested by centrifugation. The pellets were resuspended in 300 μl of 1x sample buffer (2% SDS, 10% glycerol, 125 mM Tris-HCl, pH 6.8). Zirconia beads (300 μl) were added, and cells were lysed by bead beating using a bench-top vortex mixer (60 seconds vortexing, 60 seconds on ice; 4 rounds). The homogenate was cleared by centrifugation and the protein amount measured using Pierce™ BCA protein assay (Thermo Fisher Scientific, Waltham, MA, USA). Following dilution of samples to the desired concentration, bromophenol blue (0.002%) and 2-beta-mercaptoethanol (2%) were added and samples were boiled at 95°C for 5 minutes. Proteins were resolved by SDS-PAGE and transferred to a nitrocellulose membrane using a semidry transfer apparatus. The membranes were blocked in 5% skim milk in TBST buffer (20 mM Tris-HCl (pH 7.6, 150 mM NaCl, 0.05% Tween20) for 60 min. They were then incubated overnight at 4°C in 0.05% NaN_3_-containing 1% skim milk in TBST buffer with respective primary antibodies: *Sc*Nce102 – αNce102 rabbit polyclonal (1:1000; (Loibl *et al*, 2010)); *Ca*Nce102-GFP – αGFP (1:1000; mouse): Living Colors® A.v. Monoclonal Antibody (JL-8). The membranes were washed with TBST and incubated at room temperature in the dark for 1 hour, in 1% skim milk in TBST buffer with respective secondary antibodies: αRabbit (1:20 000; goat) – LI-COR (IRDye® 800CW Goat anti-Rabbit IgG (H+L); αMouse (1:20 000; goat) – LI-COR (IRDye® 800CW Goat anti-Mouse IgG (H+L). For loading normalization, identical SDS-PAGE as described above was performed in parallel. The gels were stained in a Coomassie Blue solution (0.1% Coomassie Brilliant Blue R-250, 40% methanol, 10% glacial acetic acid) for 1 hour and destained (40% methanol, 10% glacial acetic acid) overnight on a shaker. Both the membranes and Coomassie Blue-stained gels were imaged using an Odyssey CLx infrared scanner and analysed using Image Studio Lite Ver 5.2 software (both LI-COR Biosciences, Lincoln, NE).

For the analysis of changes in *Sc*Nce102 protein abundance in response to inhibition of sphingolipid biosynthesis, exponential wild type BY4742 cultures were processed as described previously (Vaskovicova *et al*, 2020) with the following modifications: Pellets obtained by harvesting were immediately frozen in liquid nitrogen and kept at -80°C typically until the following morning when they were thawed on ice and resuspended in TNE buffer containing protease inhibitors. After diluting homogenates to desired concentration and addition of 5x Laemmli sample buffer, samples were boiled at 95°C for 5 min. Primary antibodies used: Nce102 – αNce102 rabbit polyclonal (1:1000; (Loibl *et al*, 2010)); αTubulin rat monoclonal (1:10000, ab6160, Abcam, Cambridge, UK; loading control). HRP-conjugated secondary antibodies: Nce102 – anti-rabbit; tubulin – anti-rat (goat, 1:10, 000, Jackson ImmunoResearch, West Grove, PA, USA). HRP chemiluminescence was monitored with Azure c400 (Azure Biosystems, Dublin, CA, USA) detection system and analysed using Image Studio Lite Ver 5.2. Statistical analysis was performed in GraphPad Prism 9 software.

### Lipid analysis

*S. cerevisiae* BY4742 and BY4742 *nce102*Δ cultures were grown for 48 hours as described above in the absence or presence of indicated stress conditions, washed twice with distilled water and harvested by centrifugation. The cells were counted manually using a Bürker chamber and aliquots of 5.10^8^ cells total were made. The suspensions were pelleted and C17-sphingolipids were added before cell lysis (Singh & Del Poeta, 2016; Singh *et al*, 2017). Mandala extraction was carried out as described previously (Mandala *et al*, 1995), followed by Bligh and Dyer extraction (Bligh & Dyer, 1959). A third of each sample obtained from the Bligh and Dyer extraction was reserved for inorganic phosphate (P_i_) determination. The remaining 2/3 of the organic phase were transferred to a new tube and submitted to alkaline hydrolysis of phospholipids (Clarke & Dawson, 1981). Finally, the organic phase was dried and used for lipid chromatography-mass spectrometry (LC-MS) analysis as described previously (Singh *et al*, 2017). The relative sphingolipid signal was normalized by the C17-sphingolipids and P_i_ abundance.

## DATA AVAILABILITY

All data supporting the findings reported here are available within the manuscript and its associated files. This study includes no data deposited in external repositories. Custom-written Fiji (ImageJ) macros used for microscopy image analysis are available at GitHub (https://github.com/jakubzahumensky/Nce102_SL_paper). The expanded view of this article is available online.

## ACKNOWLEDGEMENTS

We thank Izolda Mileva and Robert Rieger from the Stony Brook Cancer Center Biological Mass Spectrometry Shared Resource for expert assistance with the LC-MS analysis, and Bruce Futcher for providing us with *S. cerevisiae nce102*Δ strain used for this analysis. We further acknowledge Vendula Stradalova from Roger Schneiter lab (Université de Fribourg, Switzerland) for providing us with the Y549 *S. cerevisiae* strain.

This research was supported by the Czech Academy of Sciences, grant MSM200391901 (JZ); Czech Science Foundation, projects 19-04052S and 20-04987S (JZ, JM and PV); National Institutes of Health, grants R01AI047837 (JBK), R01AI125770 (MDP) and by a Merit Review Grant I01BX002924 (MDP) from the Veterans Affairs (VA) Program. In addition, Maurizio Del Poeta is a recipient of the Research Career Scientist (RCS) Award (IK6BX005386) and a Burroughs Welcome Investigator in Infectious Diseases.

## AUTHOR CONTRIBUTIONS

JZ and JM conceptualized and designed the study. JZ, JM, JBK and MDP secured funding. JZ and PV collected microscopy data. JZ processed microscopy data, wrote macros and scripts for their analysis and performed the analysis. JZ performed and analysed western blot analyses. JZ and CMF prepared samples for lipid analysis by mass spectrometry and analysed the data. JZ and JM wrote the manuscript, with critical input from all authors.

## EXPANDED VIEW LEGENDS

**Figure EV1.**
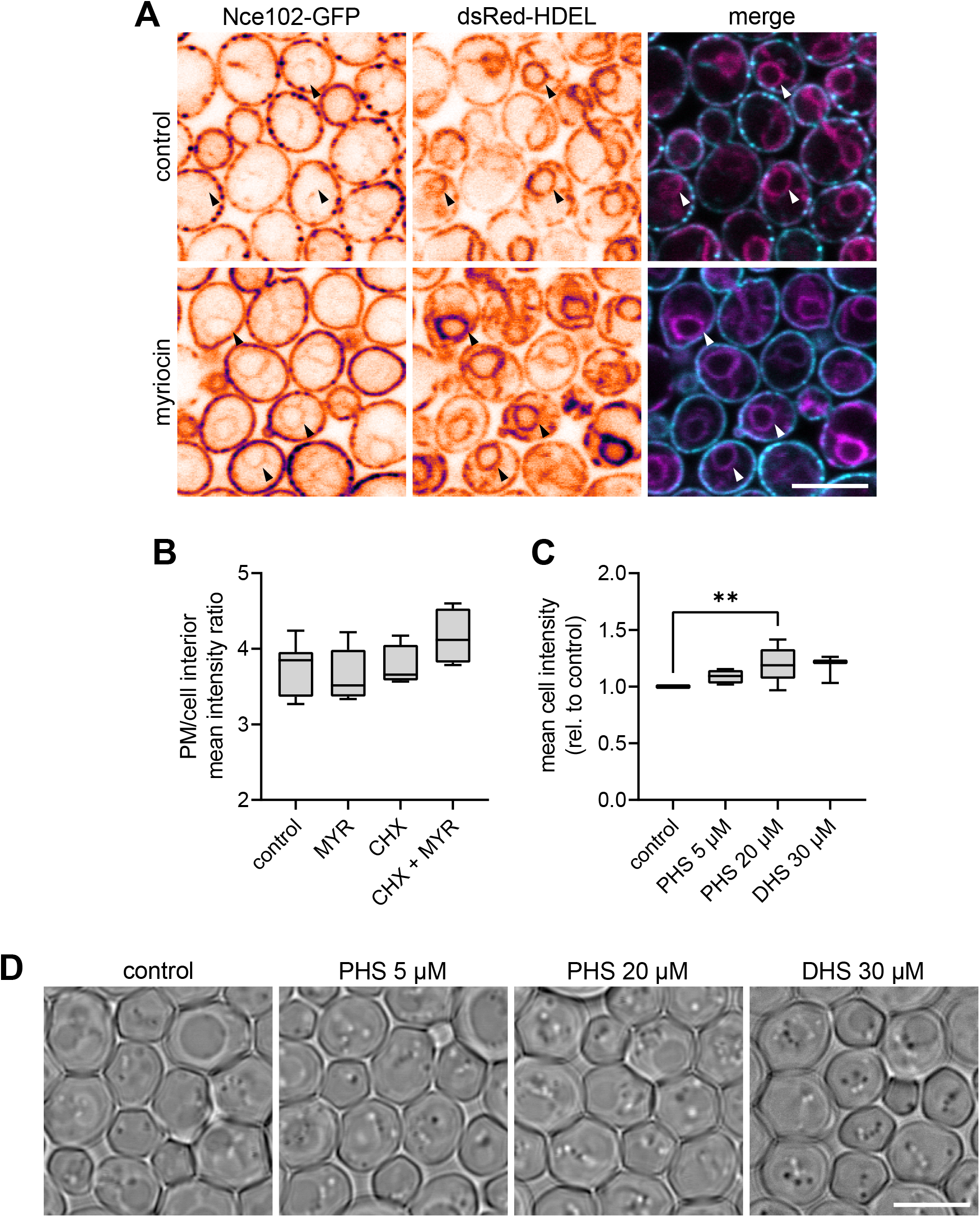
Myriocin treatment induces expression of Nce102. A Confocal microscopy images of *S. cerevisiae* cells expressing *NCE102-GFP* (cyan) together with the ER marker *dsRed-HDEL* (magenta) cultivated for 6 hours and exposed to 10 μM myriocin for 2 hours. Arrowheads point to endoplasmic reticulum. Scale bar: 5 μm. Single-channel images: LUT – inverted gem (depicted on the right-hand side of Fig 1A). B, C Quantification of the ratio of mean GFP fluorescence intensity in the plasma membrane (PM) and the cell interior (B) and mean cell GFP intensity (C) in cultures grown for 6 hours and treated with indicated stress for 2 hours. Data are presented as box plots. Bars represent the median. Error bars range from min to max value. D Bright field confocal microscopy images corresponding to Fig 1H. Scale bar: 5 μm. Data information: myriocin (MYR) – 10 μM, cycloheximide (CHX) – 100 μg/ml, DHS – dihydrosphingosine, PHS – phytosphingosine. In (B), 4-7 biological replicates (n = 170–230 cells in each condition). In (C), 3 biological replicates (n = 300–400 cells in each condition). In (C), ** – P ≤ 0.01. One-way ANOVA. No statistically significant difference between the treatments was found in (B).

**Movie EV2 – Time-lapse of the change of Nce102-GFP distribution in response to myriocin treatment**.

*S. cerevisiae* cells expressing *NCE102-GFP* were cultivated for 6 hours, treated with 10 μM myriocin and imaged in a time-lapse manner. Scale bar: 5 μm. LUT – inverted gem (depicted on the right-hand side of Fig 1A).

**Movie EV3 – Bright field channel of Movie EV2**.

*S. cerevisiae* cells expressing *NCE102-GFP* were cultivated for 6 hours, treated with 10 μM myriocin and imaged in a time-lapse manner. Scale bar: 5 μm.

**Figure EV4.**
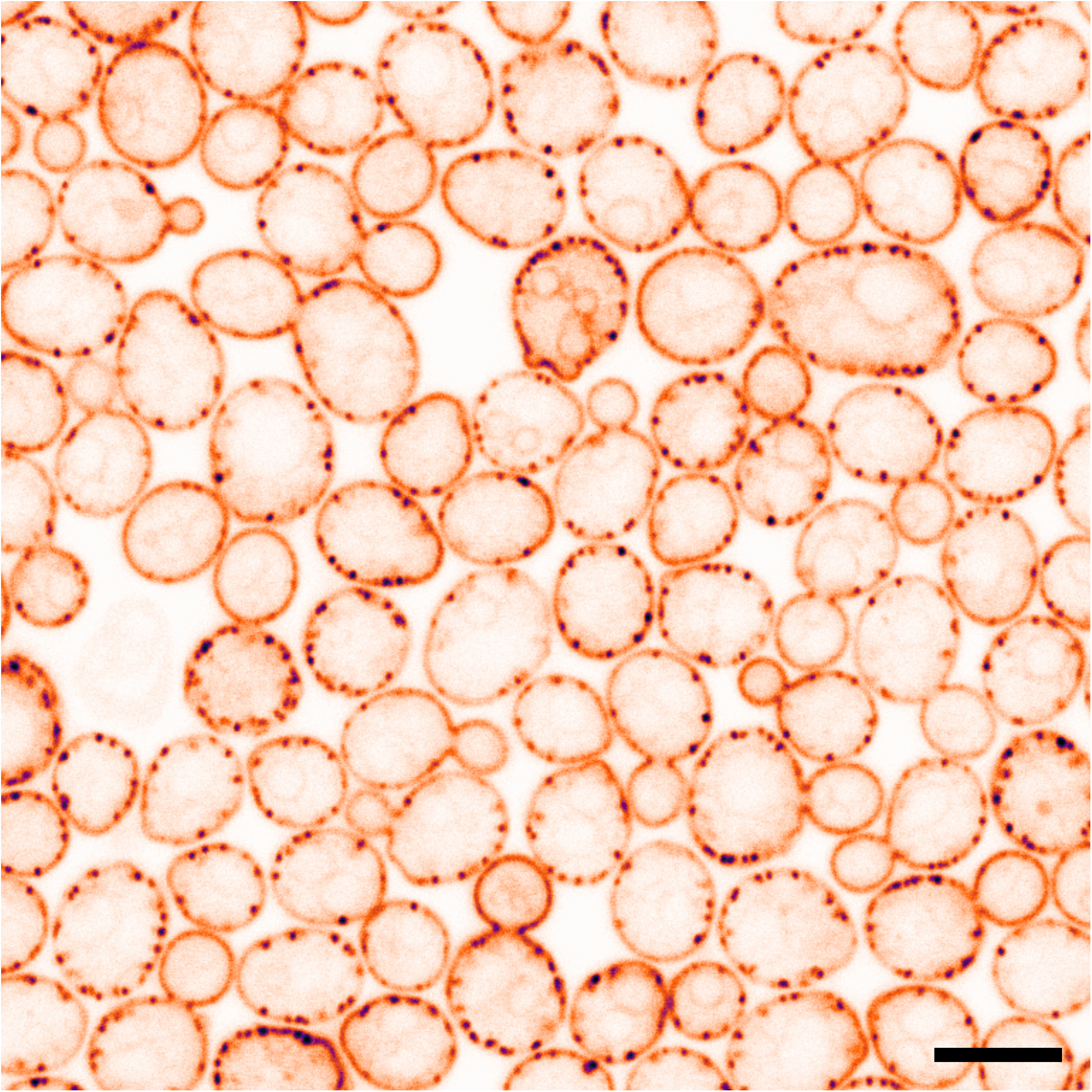
Nce102-GFP distribution in growing *S. cerevisiae* buds. Confocal microscopy image of *S. cerevisiae* cells expressing *NCE102-GFP* cultivated for 8 hours (corresponding to control sample in Fig 1A). Scale bar: 5 μm. LUT – inverted gem (depicted on the right-hand side of Fig 1A).

**Figure EV5.**
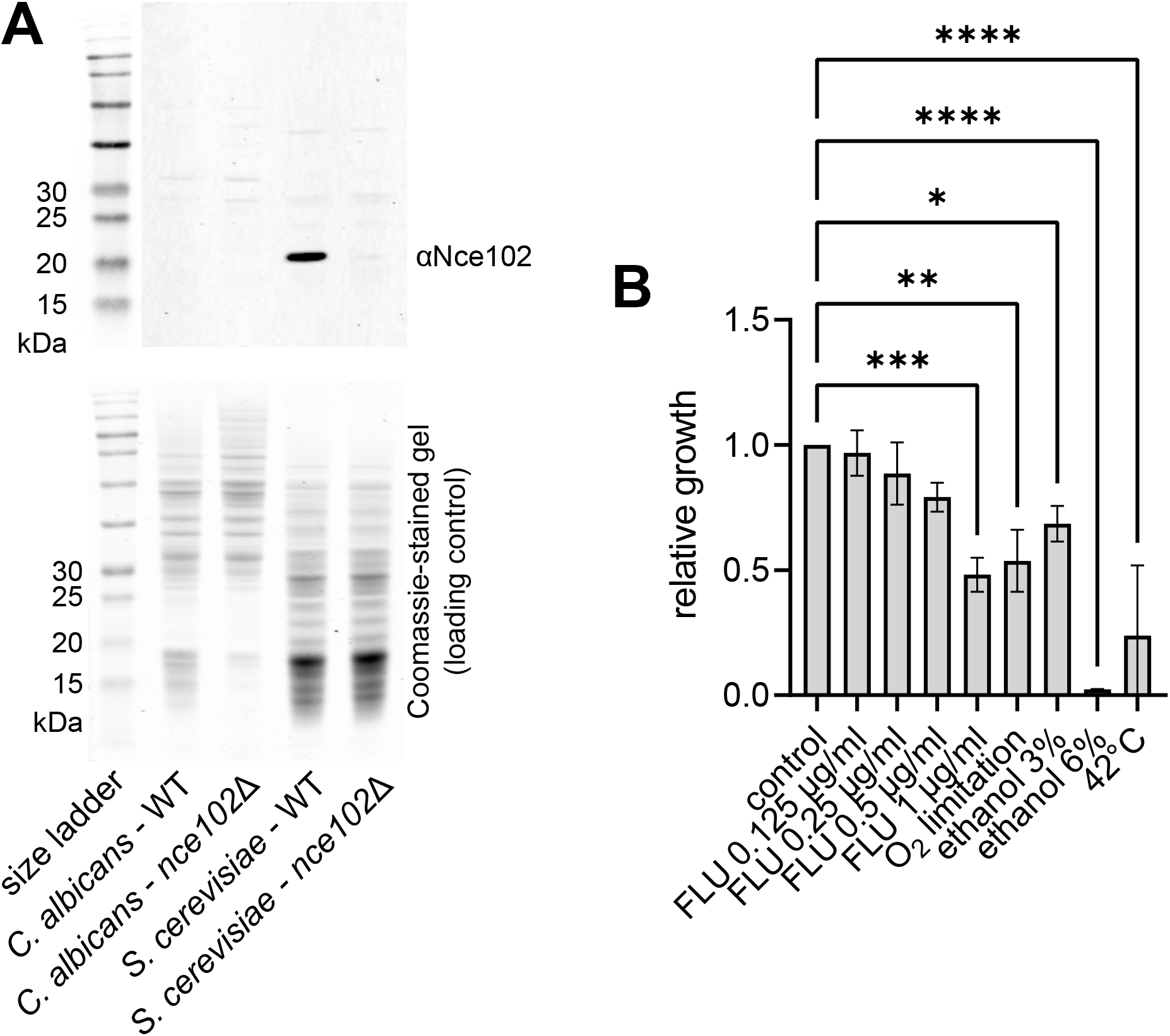
Specificity of αNce102 antibody and the effect of chronic stress on the growth of *C. albicans*. A *C. albicans* and *S. cerevisiae* wild type (WT) and *nce102*Δ cultures were grown for 24 hours and analysed by western blot for αNce102 antibody specificity. B *C. albicans* cultures were treated with indicated stress upon inoculation and cultivated for 48 hours. OD_600_ was measured and growth relative to control was calculated. FLU – fluconazole. 3 biological replicates. * – P ≤ 0.05; ** – P ≤ 0.01; *** – P ≤ 0.001; **** – P ≤ 0.0001. One-way ANOVA.

**Appendix Table S1 – Raw mass spectrometry lipid analysis data (source data for Fig 4F)**

